# *Leptospira* infection in European hedgehogs (*Erinaceus europaeus*): occurrence, diversity and associated risk factors in Greater Paris, France

**DOI:** 10.64898/2026.10.26.761796

**Authors:** Pierre-Yves Canto, Isabelle Defosseux, Emma Flandi, Mathieu Picardeau, Pascale Bourhy, Cécile le Barzic, Océane Lavaux, Julien Gasparini, Vanina Guernier-Cambert

## Abstract

**Background:** Leptospirosis is a zoonotic infectious disease whose prevalence is increasing in mainland France. While rodents are common reservoirs of pathogenic *Leptospira*, the pathogen diversity detected in French human clinical cases exceeds that currently identified in local rodents, suggesting that some unidentified host species may carry strains of public health importance. The aim of this study was to investigate *Leptospira* infection in a population of rescued injured hedgehogs (*Erinaceus europaeus*) in Île-de-France, as well as to characterize the circulating genotypes and serogroups.

**Methodology/principal findings:** In 2023-2024, kidneys from 270 dead hedgehogs collected by the Veterinary University Hospital for Wild Fauna at the National Veterinary School of Alfort, France, were tested for leptospiral DNA by real-time PCR. Twenty-two percent of individuals were qPCR-positive. Positive samples were genotyped to identify circulating species and species-groups (SGs) with *L. interrogans* SG5 being the most common. In addition, the Microscopic Agglutination Test was performed on 245 blood samples, with Australis identified as the predominant serogroup. The effect of individual and environmental variables on *Leptospira* infection status of hedgehogs was assessed, with age and season identified as significant predictors of infection.

**Conclusions/significance:** Our findings suggest that hedgehogs can harbour pathogenic *Leptospira* in Île-de-France. Further investigation is warranted to establish the epidemiological links between hedgehogs and other host species, including humans. A finer-scale analysis of the ecological features at the sampling sites could also help better understand the risk factors of *Leptospira* infection in hedgehogs at a smaller spatial scale.

## Introduction

Leptospirosis is a neglected tropical disease of global public health importance, predominantly affecting intertropical regions. Humans can become infected with pathogenic *Leptospira* species via occupational or recreational exposure to infected animals, their urine, or contaminated soil or water [1,2]. Large epidemics often occur after heavy rainfall or flooding, more commonly in tropical climates. Nonetheless, the disease is prevalent in Western Europe. An epidemiological study examining reports to the European Centre for Disease Prevention and Control from 23 European countries over the period 2010-2021 reported 12,180 confirmed leptospirosis cases overall, with France ranking first with 6,508 cases [3]. A recent report of the European Environment Agency (EEA) focusing on ‘climate change as a threat to health and well-being’ listed leptospirosis as a key climate-sensitive infectious disease [4]. Indeed, a Europe-wide spatiotemporal assessment of leptospirosis risk found that, under future climate scenarios, the incidence of leptospirosis is projected to increase [5]. In France, due to the epidemic potential of the disease, linked to its climate sensitivity and the rise of aquatic sports activities, leptospirosis has become a notifiable disease since 2023 [2]. In 2024, 871 cases of leptospirosis were reported in mainland France by the National Reference Centre, including 441 cases that were reported to the public health authorities through the mandatory notification system [6,7]. Identifying animal reservoirs of pathogenic *Leptospira* connected to human cases is key to better understanding the local epidemiological cycle of the disease. In France, rodents have been well investigated [8–11], while more sporadic studies have focused on domestic animals (cattle [11,12], swine [12,13], horses [12], dogs [12,14,15]) or wildlife (shrews [16,17], bats [18], hedgehogs [19,20], raccoons [19,21]); this is a non-exhaustive list. A study that investigated 28 animal species over 30 districts in France identified erinaceomorphs (i.e., hedgehogs) and mustelid species as having the highest risk of being infected with pathogenic *Leptospira* (OR = 66.8 and 16.7 respectively), while the highest prevalences were found in hedgehogs (37.5%) and European beavers (33.3%) [19]. Today, the exact role of urban wildlife in the maintenance and transmission of pathogenic microorganisms such as *Leptospira* to humans living in the same cities remains unclear. Features specifically associated with urban ecosystems can shape local species diversity, including both host and non-host species, which may affect both the associated pathogen diversity and the transmission and persistence dynamics within the ecosystem [22]. The Île-de-France region––which includes the city of Paris––is an urbanized environment with semi-natural spaces, parks, forests and waterways, and is host to a diverse urban fauna [23]. In the above-mentioned nationwide study focusing on 28 animal species conducted in 2012-2015, results were not provided by district, but maps were available as supporting information for erinaceomorphs and carnivores [19]. While not indicating with precision the number of animals tested, those maps showed that no *Leptospira*-positive hedgehog was identified in the Île-de-France region, whereas one carnivore (species undisclosed) tested positive. In other French districts/regions, *Leptospira*-positive hedgehogs carried predominantly *L. interrogans*, with *L. borgpetersenii* identified in a single individual. Our study aimed to explore European hedgehogs (*Erinaceus europaeus*) from the Île-de-France region as possible carriers of pathogenic *Leptospira*, focusing on a rescue population at a wildlife hospital. The objectives of our study were (1) to assess *Leptospira* qPCR positivity in this protected species, (2) to describe the *Leptospira* species, serogroup and species-group (SG) in positive individuals, and (3) to identify the individual and environmental variables (urbanization and distance to water) associated with *Leptospira* detection in hedgehogs from the Île-de-France region. Finally, we discuss our results in light of the potential role of hedgehogs as maintenance hosts for pathogenic *Leptospira* species relevant to human leptospirosis in France.

## Methods

### Sample collection

The Veterinary University Hospital for Wild Fauna (CHUV-FS) at the National Veterinary School of Alfort (ENvA), France, takes in and cares for injured, sick, or orphaned European wildlife. From April 2023 to October 2024, 1505 injured hedgehogs from the Île-de-France region were received at CHUV-FS; individuals that recovered were released in the wild, while those that died were included in our study (Fig 1). Whole carcasses were frozen at −20°C until necropsy by a veterinarian, during which their kidneys (n = 270) were collected and stored at −20°C. Individuals from this first cohort were sexed, their age determined (juvenile *versus* adult) and the date and location of their discovery before being brought to the hospital were recorded. For a subset of 185 individuals, weight was also recorded. A subset of live individuals from a second cohort (n = 131 at hospital admission, n = 76 at release) sampled between January 2023 and December 2024 were tested by serology only, using the Microscopic Agglutination Test (MAT); this cohort has been described elsewhere [24]. All procedures strictly complied with the ethical standards of the relevant French and European regulations on the care and use of animals (Directive 2010/63/EU). All procedures were approved by the ethics committee for animal experimentation (COMERC) of the National Veterinary School of Alfort: approval no. 2022-10-25, accepted 23 January 2023.

**Fig 1.**
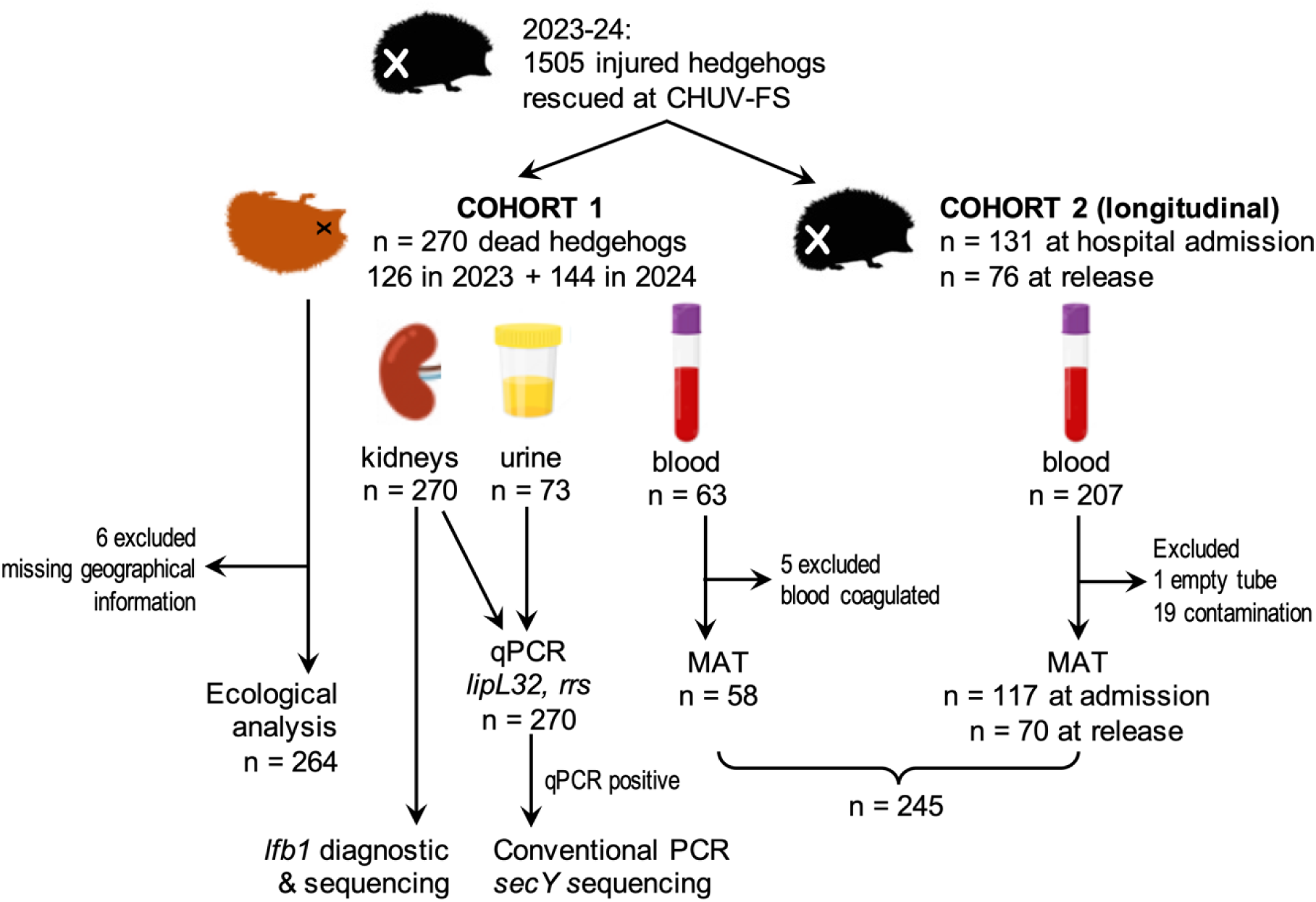
Flow chart of the study investigating hedgehogs (*Erinaceus europaeus)*. Injured animals were transported to the Veterinary University Hospital for Wild Fauna (CHUV-FS) at the National Veterinary School of Alfort (ENvA) after being rescued in the Île-de-France region. Individuals in the second cohort, which was part of a longitudinal immune study, were sampled at hospital admission and discharge to allow comparison over time (see details in [24]).

### Molecular analyses

Nucleic acid was isolated from (i) 50 mg of kidney and (ii) 200 µL of urine using High Pure PCR Template Preparation Kit® (Roche Diagnostics, Germany) according to the manufacturer’s instructions. Genomic DNA samples were stored at −20°C. Real-time PCR experiments were performed using iTaq Universal Probes One-Step® (Bio-Rad, USA) on a CFX96 Touch Real-Time PCR Detection System instrument using the following conditions: initial denaturation at 95°C for 3 min, followed by 45 cycles of denaturation for 3 sec at 95°C and annealing/elongation for 15 sec at 58°C. We used two TaqMan probes: one targeting the *lipL32* gene, which encodes a very abundant lipoprotein present only in pathogenic strains [25]; the second targeting the *rrs* gene (16S rRNA), which can detect all *Leptospira*, including non-pathogenic species [26] (S1 Table). Using a conservative approach, samples were considered positive when the cycle threshold (Ct) value was <40 in two technical replicates, keeping in mind that detection rates may be underestimated. All PCRs were run with one negative and five positive controls (*L. borgpetersenii* serovar Mini ND 2008/01925, *L. borgpetersenii* serovar Sejroe, *L. interrogans* serovar Canicola Hond Utrecht IV, *L. interrogans* serovar Icterohaemorrhagiae and *L. kirschneri* serovar Grippotyphosa reference strains).

### Microscopic Agglutination Test (MAT)

Blood was not systematically sampled, hence only a subset of individuals was tested using the MAT: 63 samples from the first cohort, some of which were also tested by qPCR, and 207 samples from the second cohort (originating from 131 hedgehogs at hospital admission; 76 hedgehogs at release), not subject to molecular testing. After exclusion of five coagulated blood samples, one empty tube, and 19 contaminations, 245 remaining samples originating from 182 individuals were tested, 58 individuals from cohort 1 and 124 from cohort 2 (Fig 1). The MAT was conducted on heparinized blood at the French National Reference Centre for leptospirosis (Institut Pasteur) using a panel of 16 antigens for the first cohort: *L. interrogans* serovar Australis, *L. interrogans* serovar Autumnalis, *L. interrogans* serovar Bataviae, *L. interrogans* serovar Canicola, *L. borgpetersenii* serovar Castellonis, *L. kirschneri* serovar Cynopteri, *L. kirschneri* serovar Grippotyphosa, *L. interrogans* serovar Hardjobovis, *L. interrogans* serovar Hebdomadis, *L. interrogans* serovar Copenhageni, *L. noguchii* serovar Panama, *L. interrogans* serovar Pomona, *L. interrogans* serovar Pyrogenes, *L. borgpetersenii* serovar Sejroë, *L. borgpetersenii* serovar Tarassovi and *L. interrogans* serovar Icterohaemorrhagiae. A reduced panel of eight antigens was used for the second cohort: *L. interrogans* serovar Australis, *L. interrogans* serovar Bataviae, *L. kirschneri* serovar Grippotyphosa, *L. interrogans* serovar Hebdomadis, *L. interrogans* serovar Copenhageni, *L. interrogans* serovar Pomona, *L. borgpetersenii* serovar Sejroë and *L. interrogans* serovar Icterohaemorrhagiae. The reduced panel was selected based on the serological reactivity observed in the first cohort. The presumptive infecting serogroup was assigned based on the serovar showing the highest titre. The MAT was considered positive when the titre was ≥ 1:200 for at least one antigen.

### Leptospira genotyping

*Leptospira* qPCR-positive samples were subjected to conventional PCR amplification using a published protocol [27] targeting the *secY* gene (202 bp fragment). All PCRs were run with negative and positive controls. PCR amplifications were performed in 50 µl containing 1 µl of nucleic acids template, 5 µl of 10x PCR Buffer mix, 1.25 µl (10 µM) of each primer, 1 µl of dNTPs (10 mM), 1.5 µl of MgCl2 (25 mM), 0.25 µl of Taq polymerase (5 U/µl; Thermo Fisher) and 38.75 µl of RNase-free water, using the following conditions: 5 min of initial denaturation at 95°C and 40 cycles of denaturation for 15 sec at 95°C, annealing for 30 sec at 58°C, extension for 30 sec at 72°C and one cycle of final extension for 5 min at 72°C using a conventional thermal cycler. PCR products were run on a 1.5% gel at 100 V for 30 min, and amplicons matching the expected size were sent to Eurofins for Sanger sequencing.

The *lfb1* genotyping method was also applied to all DNA samples at the French National Reference Centre for Leptospirosis, Institut Pasteur, using a published protocol [14]. The *lfb1* 331-bp fragment was amplified and sequenced by Sanger sequencing (Eurofins).

Primer details are provided in S1 Table.

### Phylogenetic analysis

For *secY*, consensus sequences were generated and aligned with Geneious Prime 2025 (http://www.geneious.com). For *lfb1*, sequence and phylogenetic tree analyses were performed using the software BioNumerics V7.6 (Applied-Maths, Saint-Martens-Latem, Belgium) as previously described [14]. *lfb1* sequencing was used to classify samples into *Leptospira* species-groups (SGs) based on sequence alignment against a database of *lfb1* sequences, as previously published [14].

### Environmental data

Maps were created using the open-source software QGIS 3.34.4 [28]. For each hedgehog, we calculated the sampling location. If an exact address was provided by the person rescuing the animal, the GPS coordinates were retrieved using Google Maps, taking into account details such as “found in the garden” or “in front of the garage” when specified. If only the street/avenue/park/city were provided, we used coordinates automatically generated by Google Maps.

For each hedgehog, we also quantified (1) the degree of urbanization and (2) the distance to the nearest water body. The degree of urbanization was defined as the proportion of land occupied by urban infrastructure that humans frequently use, such as buildings and roads, as previously described [29]. First, we imported the 2017 Land Use and Land Cover (LULC) available from the public database of the region Île-de-France (https://www.data.gouv.fr/datasets/mode-doccupation-du-sol-mos-en-11-postes-en-2017), along with the coordinates of the hedgehog collection points. Both layers were converted into the coordinate reference system (CRS) RGF93-Lambert 93. Second, we developed a model to estimate the percentage of urbanization by creating a 500-meter buffer zone around the collection sites in order to cover the hedgehog’s home range [30]. The sampling locations encompassed areas with heterogeneous levels of urbanization. The distance (m) to the nearest permanent water body (lake, pond or river) was calculated using the “Shortest line between features” tool in QGIS. Environmental data are detailed in S2 Table; GPS coordinates are not provided to ensure anonymity.

### Statistical analyses

To assess the association between individual and environmental factors and *Leptospira* infection status, we fitted a generalized linear model (GLM) with a binomial distribution, using infection status as the binary response variable (presence = 1; absence = 0). Age, sex, season, percentage of urbanization and distance to the nearest water body were included as explanatory variables. To avoid collinearity, body mass was not included in the model because it was correlated with sex and age. All statistical analyses were conducted using JMP®19.

## Results

### Population description

In total, 1505 injured hedgehogs rescued from all departments of the Île-de-France region were received at CHUV-FS during the study period. Of these, 270 died despite treatment and were included in our study (first cohort): 126 individuals collected in 2023 and 144 in 2024 (S2 Table). Kidney samples were collected from all 270 individuals; 73 urine samples and 63 blood samples were also available. All individuals were tested molecularly, but six individuals with missing geographical location data were excluded from ecological analyses. Spatial distribution was heterogeneous (Fig 2). The distribution by age, sex and season is provided in Table 1. The population was broadly balanced by sex and age, with a higher proportion of individuals collected during summer (42%) (Table 1). Consistent with the hibernation behaviour of this species, few individuals (3%) were collected during winter. The second cohort, described elsewhere [24], was only included in the serology testing (MAT) and not in the ecological analysis.

**Fig 2.**
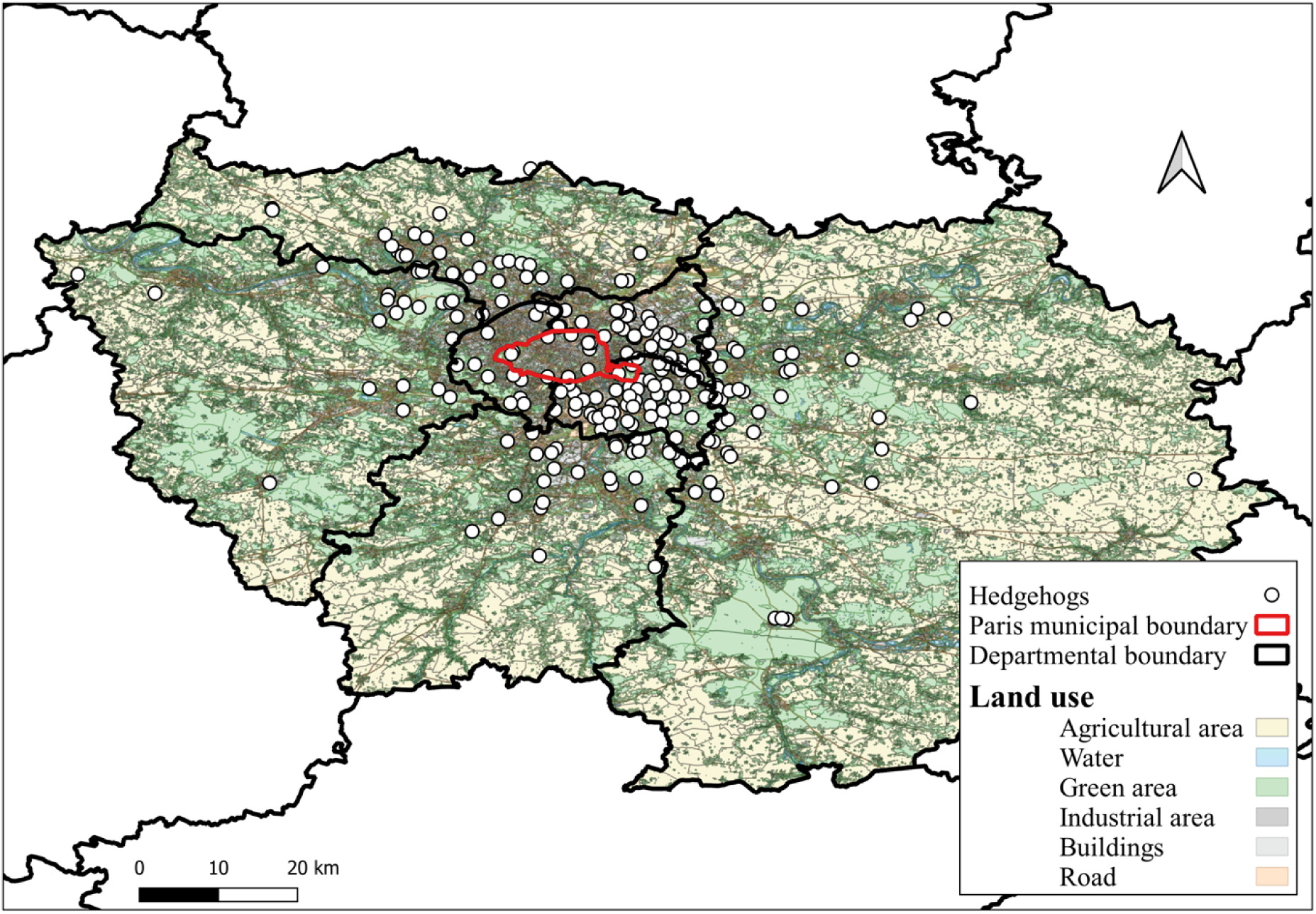
Identification of the study site (Île-de-France), land use and collection points of hedgehogs (*Erinaceus europaeus)*. Background layer is the 2017 Land Use Land Cover (LULC) available from https://www.data.gouv.fr/datasets/mode-doccupation-du-sol-mos-en-11-postes-en-2017. The map was generated using QGIS 3.34.4.

**Table 1.** Characteristics of the study population of European hedgehogs (*Erinaceus europaeus)* (n = 270) from Île-de-France (greater Paris) and distribution of *Leptospira* qPCR-positive results. Sex, age and season at the time of arrival at the hospital (CHUV-FS) were recorded. The diagnostic qPCR targeted the *lipL32* gene [25] and the *rrs* gene (16S rRNA) [26]. NA: not applicable. Six individuals with missing geographical data were later excluded from the ecological analysis.

| Hedgehogs' characteristics | n (%) | *Leptospira* qPCR-positive (%) |
| --- | --- | --- |
| **Sex** |  |  |
| Male | 134 (50%) | 31 (23%) |
| Female | 128 (47%) | 25 (19%) |
| Missing data | 8 (3%) | 3 (NA) |
| **Age** |  |  |
| Adult | 151 (56%) | 48 (32%) |
| Juvenile | 118 (44%) | 10 (8%) |
| Missing data | 1 (NA) | 1 (NA) |
| **Season** |  |  |
| Spring | 70 (26%) | 28 (41%) |
| Summer | 115 (42%) | 25 (22%) |
| Autumn | 75 (28%) | 4 (5%) |
| Winter | 9 (3%) | 1 (11%) |
| Missing data | 1 (NA) | 1 (NA) |
|  | 270 (100%) | 59 (22%) |

### Leptospira infection and diversity

A subset of urine samples yielded the same *Leptospira* qPCR results as the corresponding kidney samples from the same individuals and were therefore excluded from further analysis. Among the 270 hedgehogs tested, 59 (21.9%) were *Leptospira* qPCR-positive. Detailed Ct values obtained for each positive qPCR are provided in S2 Table. Out of these 59 *Leptospira* qPCR-positive individuals, 30 were successfully amplified by PCR with the *secY* target, and 24 yielded sequences, all of which were identified as *L. interrogans*. For the *lfb1* target, 16 samples were successfully sequenced and identified as *L. interrogans* (SG1 and SG5) and *L. kirschneri* (SG4) (Table 2). Within the group of identified SGs, *L. interrogans* SG5 was the most commonly identified (13/16, 81%), followed by *L. interrogans* SG1 (2/16, 13%) and *L. kirschneri* SG4 (1/16, 6%). GenBank accession numbers of the sequences are provided as supplementary information (S3 Table).

**Table 2.**
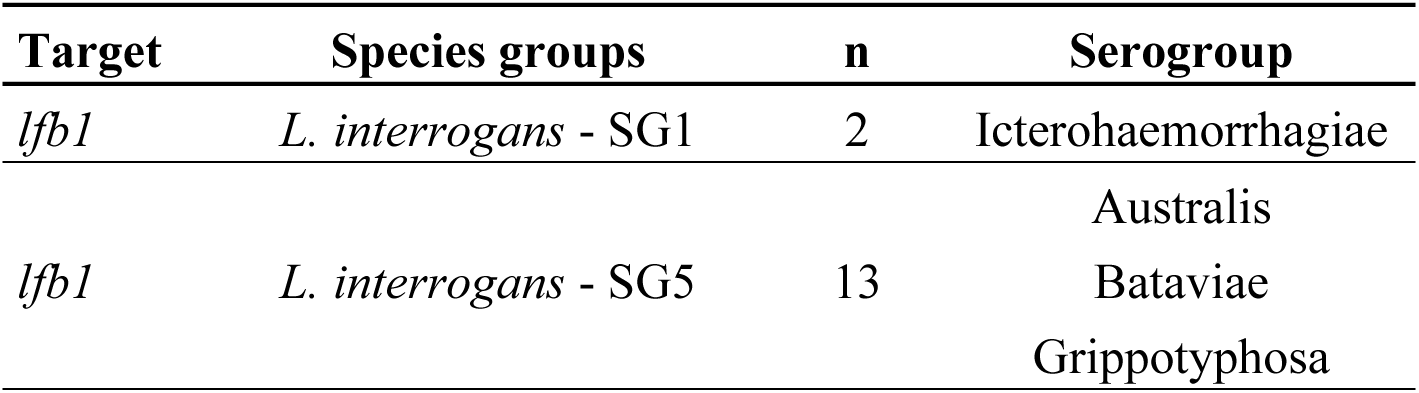
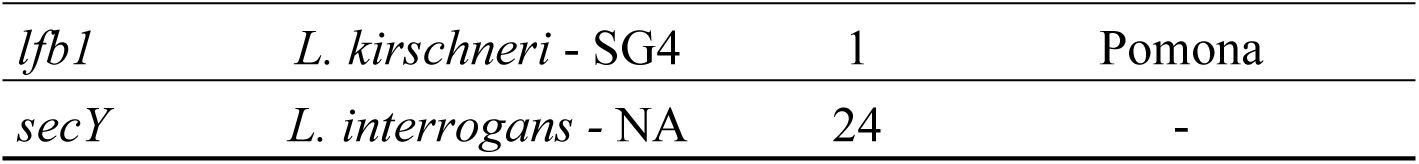
Diversity analysis of *Leptospira*-positive hedgehogs based on *lfb1* (n = 16) and *secY* sequences (n = 24). *Leptospira* species groups (SGs) are defined on the basis of *lfb1* sequence polymorphism. n: number of individuals in each species group. NA: not applicable. Presumptive serogroup(s) associated with each SG, based on previous characterization [14], are provided.

### MAT results

After the exclusion of five coagulated samples, 58 samples from the first cohort were tested with 16 antigens, and 25 samples had a positive MAT reaction at a dilution of ≥ 1:200. Of these, 23 were seroreactive to a single serovar: 22 to sv Australis and one to sv Pomona. Two samples were seroreactive to three serovars: one sample to sv Australis, Grippotyphosa and Panama, and one sample to sv Autumnalis, Grippotyphosa and Pomona (S4 Table).

After exclusion of one empty tube and 19 contaminations, 187 blood samples from the second cohort, including 63 paired samples, were tested with 8 antigens; 39 samples had a positive MAT reaction at a dilution of ≥ 1:200, including 25 non-redundant results. All were seroreactive to a single serovar: 38 to sv Australis and one to sv Grippotyphosa. Five had MAT titres ≥ 1:1600. To our knowledge, this was the first time MAT was attempted on heparinized blood. Detailed MAT results of seroreactive individuals (at hospital admission and at release for paired samples) are provided as supplementary information (S4 Table). Among individuals for whom both molecular and serological results were available, three were both qPCR-positive and seroreactive; all three were classified as *L. interrogans* SG5, corresponding to a presumptive Australis serogroup, which was confirmed by the MAT.

### Impact of individual and environmental factors on Leptospira infection status

Our ecological analysis included 264 individuals after excluding six individuals with missing geographical data, all *Leptospira* qPCR-positive. The statistical model identified significant associations between *Leptospira* infection status and age and season, whereas no significant associations were detected with sex, percentage of urbanization, or distance to nearest water body (Table 3). First, the proportion of qPCR-positive individuals was higher among adults (30%; 45/148) than among juveniles (7%; 8/116). Second, the proportion of qPCR-positive individuals also varied significantly among seasons, with higher proportions observed in spring and summer than in autumn and winter.

**Table 3.**
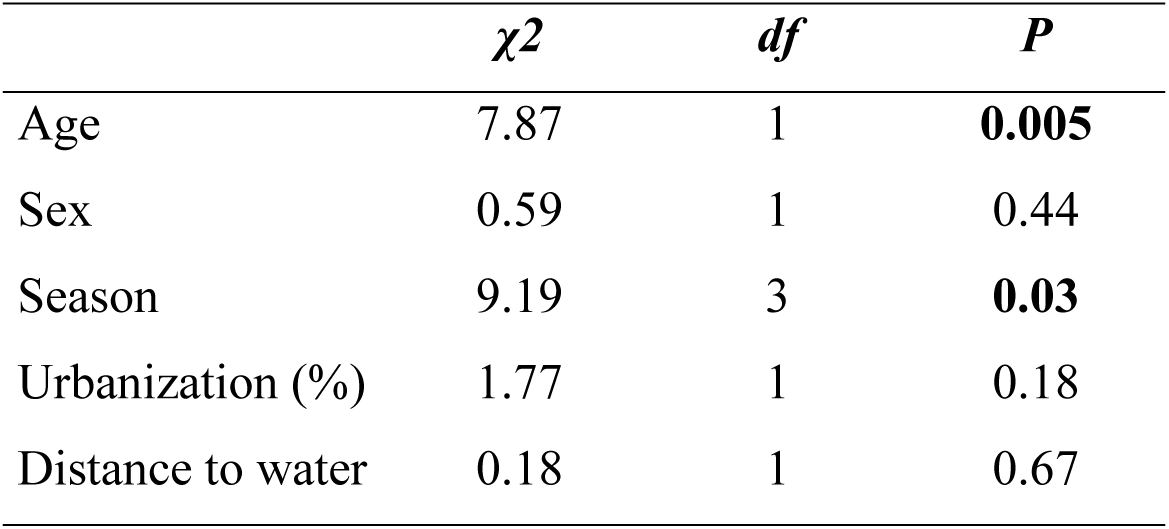
Results of the GLM assessing associations between age, sex, season, percentage of urbanization and distance to nearest water body and *Leptospira* infection status of hedgehogs (n = 264). Infection status is *Leptospira* qPCR-positive versus *Leptospira* qPCR-negative. χ2: results from likelihood-ratio χ² tests. *df*: degrees of freedom. Statistically significant associations (P < 0.05) are in bold.

| | $\chi^2$ | *df* | *P* |
| --- | --- | --- | --- |
| Age | 7.87 | 1 | **0.005** |
| Sex | 0.59 | 1 | 0.44 |
| Season | 9.19 | 3 | **0.03** |
| Urbanization (%) | 1.77 | 1 | 0.18 |
| Distance to water | 0.18 | 1 | 0.67 |

## Discussion

This study investigated *Leptospira* infection in an injured rescued hedgehog population in Île-de-France by PCR and serology. Our molecular results indicate that pathogenic *Leptospira* occur in hedgehogs in Île-de-France, with 22% (59/270) of the study population testing positive. This is lower than the 37.5% (42/112) prevalence reported in hedgehogs at the national level in France in 2012-2015 [19] or the 36% (25/69) urinary prevalence in rescued hedgehogs in Nantes in 2022 [20]. Those differences could reflect regional or temporal variations in France or could result from sampling bias, small sample sizes or laboratory procedures. By comparison, a study conducted in North-western Italy reported a 17.6% prevalence in hedgehog carcasses collected by the Public Veterinary Service, which is more similar to our results [31]. In the Île-de-France specifically, previous investigations of *Leptospira* infections were limited to rats. A study investigating brown rats in Paris identified pathogenic *Leptospira* in 15% of individuals, all carrying *L. interrogans* [32].

Among the samples successfully characterized by sequencing, *L. interrogans* was predominant, with *L. kirschneri* identified in one individual. *L. interrogans* is commonly detected in French wild mammals. In a national study conducted in France in 2012-2015, *L. interrogans* (n = 140) was identified in all five groups of wild animals, while *L. kirschneri* (n = 37) and *L. borgpetersenii* (n = 25) were detected in large carnivores, mustelids and ungulates but not in lagomorphs and rodents, though the authors did not provide the details per species [19]. In this same study, one hedgehog was infected with *L. borgpetersenii*, while the others (n = 41) carried *L. interrogans.* In a 2020-2022 study, *L. kirschneri* was detected in three greater white-toothed shrew (*Crocidura russula*) in Lyon, France [16]., Together with previous studies, our results indicate that, although *L. interrogans* is the dominant species overall, *L. kirschneri* and *L. borgpetersenii* have also been detected in hedgehogs in France.

In terms of species-groups, among the 16 individuals successfully characterized in our study, SG5 was predominant, being identified in 13 individuals. In a 2019-2021 study in mainland France, *L. interrogans* SG1 was reported to be the most commonly (53/110 = 48%) identified in humans, followed by *L. interrogans* SG5 (9/110 = 8%). The pattern was similar in dogs, though with different frequencies: *L. interrogans* SG1 was the most frequently detected (65%; 39/60) followed by *L. interrogans* SG5 (25%; 15/60) [14]. However, a study conducted in dogs in 2020-2024 showed that *L. interrogans* SG5 was predominant (47.3%), followed by *L. interrogans* SG1 (45.6%) [33]. These findings illustrate variation in the relative frequency of SGs among host populations and study periods.

Previous studies have shown that *L. interrogans* SG5 primarily corresponds to serogroup Australis [14,33]. This is coherent with our MAT serology results, with reactivity to Australis observed in 61 of 64 seroreactive samples. Interestingly, Australis was already identified in French hedgehogs collected in 1960 in Normandie, and in 1973 in Maine-et-Loire, Ille-et-Vilaine and Loire-Atlantique [34]. More recently, Australis has also been detected among human patients in France [6], although less frequently than in dogs and hedgehogs.

In the hedgehog population of Île-de-France, the proportion of qPCR-positive individuals was highest in spring (40%, 28/70), followed by summer (22%, 25/115), winter (11%, 1/9) and autumn (5%, 4/75). Season was significantly associated with *Leptospira* infection status in the GLM. In the Italian study focusing on several animal species in the Po Valley, the hedgehog was the only species showing a statistically significant seasonality of *Leptospira* infection, with the highest prevalence in summer (25%) and the lowest prevalence in winter (8%) [31]. The highest proportion of positive individuals in spring observed in our study may reflect changes in host condition or immune status following winter hibernation [35] while the lowest proportion in winter should be interpreted with caution because of the small sample size during this season. The seasonality pattern was different in Italy, which may be related to a different risk factor; the authors of the study suggested that yearly rainfall may impact seasonality patterns. Wildlife rehabilitation centres, although they provide access to a large sample size of a protected species, generate cohorts of individuals that are non-representative of the wider population, both with respect to animal health status and timing of admission. Nevertheless, such samples can provide a valuable first approach for identifying broad patterns [29], which should subsequently be confirmed in representative populations.

Our results suggest that *Leptospira* infection is more frequently detected in adult than juvenile hedgehogs. This pattern has been observed in different rodent host species, and has been attributed to increased probability of exposure as well as cumulative exposure with age [36]. Leptospirosis has often been discussed as a water-borne disease and water plays an important role in the environmental persistence of *Leptospira* [37,38]. Consequently, proximity to water has been examined as a potential factor associated with *Leptospira* infection. In the Republic of Sakha, Russian Federation, the likelihood of infection in livestock and wild animals decreased as the distance to water increased [39]. In Lithuania, *Leptospira* prevalence in striped field mice and yellow-necked mice was significantly negatively correlated with the distance to water [40]. In Kansas and Nebraska, USA, distance to water (OR = 0.8), hydrologic density (OR = 2.8) and frequently flooded areas (OR = 4.1) within 2500 m surrounding case/control locations were significant risk factors for canine leptospirosis [41]. However, distance to water was not significantly associated with *Leptospira* infection status in hedgehogs in our study. This may be related to regional features specific to Île-de-France: in this highly urbanized environment, small water sources such as puddles, backyard ponds, fountains, or water containers provided by people for wildlife and/or domestic animals, could contribute to *Leptospira* exposure in hedgehogs and may not have been captured by our GIS-based analysis which focused on larger permanent water bodies.

Altogether, our findings suggest that hedgehogs are a reservoir of pathogenic *Leptospira* in Île-de-France. In our study population, we identified *Leptospira* DNA in 22% of rescued hedgehogs, and *L. interrogans* SG5 and serogroup Australis predominated among the molecularly and serologically characterized samples, respectively. These genotype and serogroup are also prevalent in humans and dogs in France. While dogs are susceptible to infections with Australis, hedgehogs appear unaffected, suggesting co-adaptation and a role as healthy chronic carriers, similar to rats and mice with Icterohaemorrhagiae. This is supported by a previous study demonstrating that infected hedgehogs exhibit chronic *Leptospira* shedding, with no body weight loss, and maintain infection even after antibiotic treatment [20]. Establishing the epidemiological links between hedgehogs and other host species, including humans, warrants further investigation. Finer-scale characterization of environmental features at collection sites, including small water sources, could also help identify ecological factors associated with *Leptospira* exposure in hedgehogs.

## Acknowledgements

We thank all the volunteer citizens that rescued the animals and brought them to the Veterinary University Hospital for Wild Fauna (CHUV-FS) at the National Veterinary School of Alfort (ENvA). We express our gratitude to the NGO Faune Alfort for its constant support of the work of CHUV-FS (Veterinary University Hospital for Wild Fauna) and for granting us permission to conduct our research within their facilities. We are also grateful to the entire care team— Camille Choutet, Elyssa Della, Inès de Sousa, Noémie Raphaël, Nora Affejee, Valentin Delon, Zoé Warren—and to all volunteers for their invaluable assistance. We also thank the staff of the French National Reference Centre for Leptospirosis (technicians Jean-François Mariet, Faida Zinini and Céline Lorioux) for performing the *lfb1* genotyping and MAT.

